# Assessing the role of *At*GRP7 arginine 141, a target of dimethylation by PRMT5, in flowering time control

**DOI:** 10.1101/2024.09.24.614656

**Authors:** Alexander Steffen, Katarzyna Dombert, María José Iglesias, Christine Nolte, María José de Leone, Marcelo J. Yanovsky, Julieta L. Mateos, Dorothee Staiger

## Abstract

Arginine (R) methylation, catalyzed by PROTEIN ARGININE METHYLTRANSFERASES (PRMTs), is critical for regulation of gene expression at the transcriptional and post-transcriptional level. Among nine PRMT genes in Arabidopsis, PRMT5 catalyzing symmetric R dimethylation of its targets is best characterized. PRMT5 mutants are late flowering and show altered responses to environmental stress. Among PRMT5 targets are *Arabidopsis thaliana* GLYCINE RICH RNA BINDING PROTEIN 7 (*At*GRP7) and *At*GRP8 that promote the transition to flowering. *At*GRP7 R141 has been shown to be modified by PRMT5. Here, we tested whether this symmetric dimethylation of R141 is important for *At*GRP7’s physiological role in flowering time control. We constructed *At*GRP7 mutant variants with non-methylable R141 (R141A, R141K). Genomic clones containing these variants complemented the late flowering phenotype of the *grp7-1* mutant to the same extend as wild-type *At*GRP7. Furthermore, overexpression of *At*GRP7 R141A or R141K promoted flowering similar to overexpression of the wild-type protein. Thus, flowering time does not depend on R141 and its modification. However, *At*GRP7 R141 contributes to the activity of GRP7 in response to abscisic. Immunoprecipitation of *At*GRP7-GFP in the *prmt5* background revealed that antibodies against dimethylated arginine still recognized *At*GRP7, suggesting that additional methyltransferases may be responsible for modification of *At*GRP7.

## 1. Introduction

Modification of arginine (R) residues, although discovered half a century ago, has only recently been recognized to play a key role in the regulation of transcription, posttranscriptional control and DNA repair (1). R methylation is catalyzed by protein arginine methyl transferases (PRMTs) with S-Adenosyl-methionine as a methyl donor. The importance of correct R methylation is underscored by impaired PRMT activity associated with autoimmune diseases or cancer in mammals (2). *Arabidopsis thaliana* contains nine PRMT genes (3). Best characterized is PRMT5, also known as Shk1 binding protein 1 (SKB1), a type II PRMT which catalyzes the formation of symmetric dimethylarginine. *Prmt5* mutants show a variety of defects including late flowering, reduced sensitivity to vernalization, a long period circadian phenotype and reduced sensitivity to salt stress (4-8).

*At*PRMT5 was shown to add two methyl groups to arginine three of histone H4 to form H4R3sme2, which is a repressive mark for gene transcription (9). It has been proposed that PRMT5 methylates H4R3 in the promoter of the key floral repressor FLOWERING LOCUS C (FLC). This leads to suppression of

*FLC* expression and flower induction (4).

The use of antibodies that detect symmetrically dimethylated arginines such as the SYM10 antibody, raised against a peptide containing four symmetrical dimethyl-arginine-glycine repeats, allowed the detection of a wide spectrum of substrates in wild type (wt) plants, but not in the *prmt5* mutant (10). Among those were, in addition to histones, core spliceosomal U small nuclear ribonucleoproteins including *At*SmD1, *At*SmD3 and *At*LSm4 (10,11). In mammals, methylation increases the binding affinity of the Sm proteins to SURVIVAL OF MOTOR NEURON (SMN) to promote assembly of the spliceosome (12,13). Loss of symmetric arginine dimethylation of Sm proteins in the Arabidopsis *prmt5* mutant has recently been shown to prevent the recruitment of the Nineteen complex to the spliceosome and initiation of spliceosome activation (14). Recently, we showed that methylation of AtLSm4 fine-tunes splicing in response to stress (15).

Indeed, global splicing defects were observed in the *prmt5* mutant (6,10,16). In *prmt5*, the transcript encoding FLK, a component of the autonomous pathway of flowering time control, is misspliced. An elevated level of unproductive transcript with retained intron 1 is observed at the expense of the transcript encoding the functional protein, which contributes to the late flowering of *prmt5* (14).

Among the PRMT5 substrates are also numerous proteins involved in RNA processing including the circadian clock regulated *At*GRP7 (*A. thaliana* glycine-rich RNA-binding protein 7) and *At*GRP8 proteins (10). These are similar to mammalian hnRNP like proteins and consist of an N-terminal RNA recognition motif (RRM) and a C-terminal glycine-rich stretch.

*At*GRP7 is part of the circadian timing system, promotes the transition to flowering and the defence against pathogenic bacteria. Furthermore, *At*GRP7 exhibits RNA chaperone and nuclear export function and promotes freezing tolerance (17,18). *In vivo* targets of *At*GRP7 have been determined by individual nucleotide resolution UV cross-linking and immunoprecipitation (iCLIP) (19,20). *At*GRP7 affects splicing of some of its targets as well as processing of a suite of miRNA precursors (21). The conserved R49 in the RRM is crucial for *in vivo* binding activity and function. Furthermore, truncation or deletion of the glycine-rich C-terminal domain reduced *in vitro* binding (22,23). For *At*GRP7 and *At*GRP8, R141 located in the glycine-rich domain was identified by mass spectrometry to be the residue methylated *in vivo* by PRMT5 but data on the physiological consequences of this modification are so far lacking (10). Here, we set out to determine the relevance of R141 dimethylation for its function in flowering time.

We mutated the R141 residue so that it can no longer be methylated. Genomic constructs expressing the nonmethylable *At*GRP7 complemented the late flowering phenotype of the *grp7-1* mutant to the same extent as wt *AtGRP7*. Furthermore, constitutive overexpression of nonmethylable *At*GRP7 in wt plants promoted flowering to the same extent as plants overexpressing the authentic proteins. In contrast, physiological experiments with abscisic acid (ABA) showed that R141 methylation may be important under stress. To determine the methylation status, we immunoprecipitated *At*GRP7-GREEN FLUORESCENT PROTEIN (GFP) from wt and *prmt5* backgrounds and found that dimethylation of *At*GRP7-GFP was reduced but not abolished, raising the possibility that PRMT5 might not be the sole methyltransferase responsible for *At*GRP7 modification.

### 2. Results

### 2.1. Genomic *At*GRP7 R141 variants complement late flowering of *grp7-1*

The *grp7-1* mutant flowers with a higher leaf number than wt type plants particularly in SDs (24,25). To test whether R141 is critical for the floral promoting effect of *At*GRP7, we mutated R141 to alanine or lysine in the *At*GRP7 genomic clone and introduced the wt and mutated genomic constructs into the *grp7-1* mutant. Whereas *grp7-1* flowered with more leaves than Col-0 plants, two independent lines complemented with the wt type genomic construct (FL4a (full length 4a) and FL10c) flowered with a similar leaf number as Col-0 plants (Fig. 1a). Four independent lines expressing the GRP7 R141A variant (lines 3, 4, 8, 13) also flowered with similar leaf numbers as Col-0. From five independent lines expressing the GRP7 R141K mutation, again four lines flowered similar to Col-0 plants whereas one transgenic line (#5) flowered like *grp7-1* (Fig. 1a). We monitored *At*GRP7 protein abundance in the different transgenic lines and found variable expression levels in most independent lines (Fig. 1b). Notably, GRP7 R141K line 5 which flowered with the highest leaf number had the lowest *At*GRP7 level (Fig. 1b). This suggests that flowering time is independent of whether it contains R141 or mutations thereof. Abundance of the closely related *At*GRP8 was elevated in *grp7-1* due to relief of repression by *At*GRP7 (26) and below wt level in FL4a and FL10c (Fig. 1b). In GRP7 R141A lines the *At*GRP8 level was low in line 4 that expressed more *At*GRP7 than wt and high in *grp7-1* and all complemented lines with low *At*GRP7 expression (Fig. 1b), leading us to conclude that the negative regulation of *AtGRP8* through *At*GRP7 is still intact.

**Figure 1.**
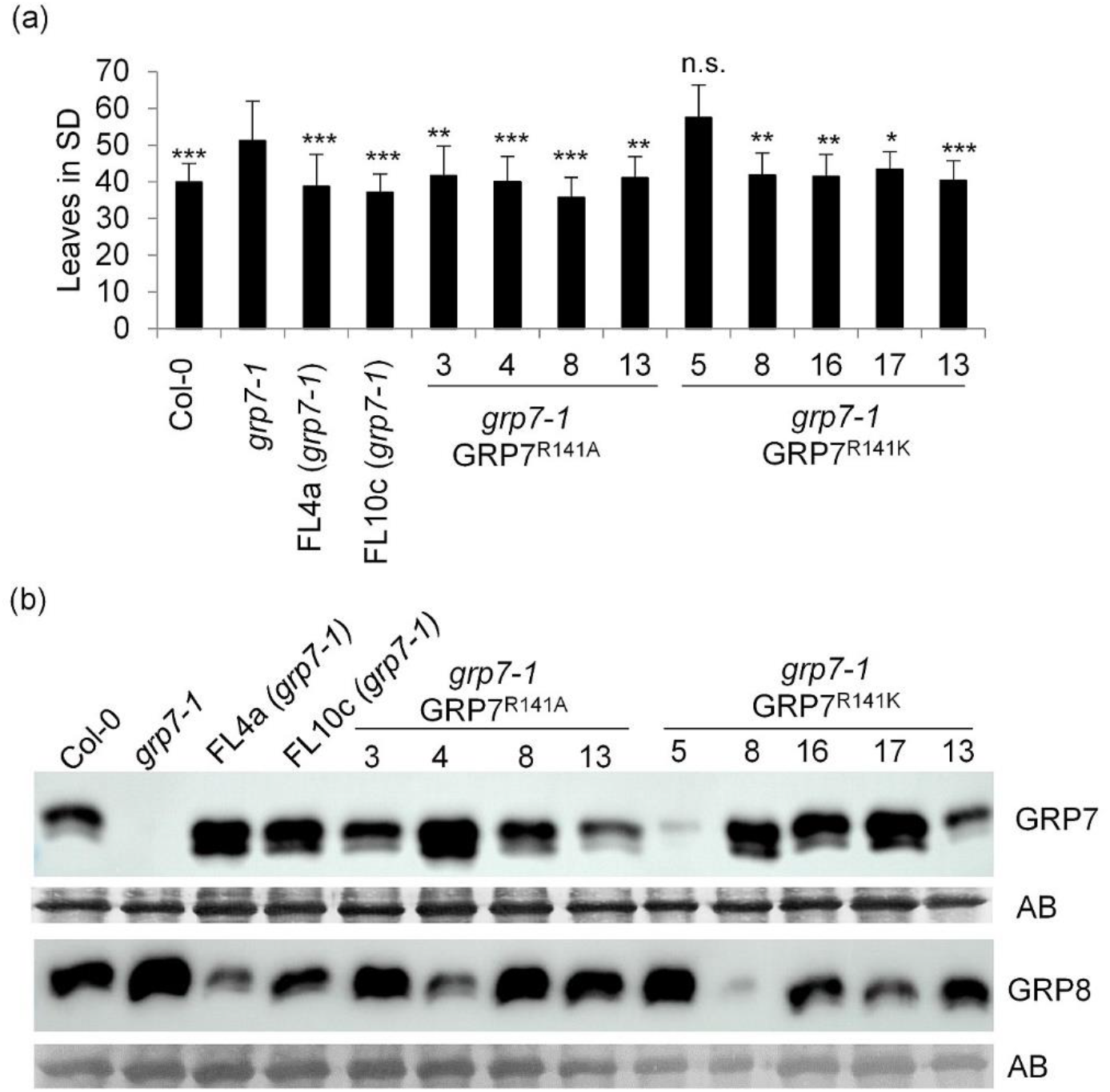
Flowering time of the *grp7-1* mutant complemented with genomic *At*GRP7 and R141 variants. (a) Col wt, *grp7-1*, two *grp7-1* lines complemented with genomic *At*GRP7 (FL4a and FL10c), *grp7-1* lines complemented with genomic *At*GRP7 R141A or genomic *At*GRP7 R141K were grown in SDs (n = 15-20). The number of rosette leaves are shown as mean ± SD. ANOVA followed by a Dunnett’s test was performed to determine statistical significance (* P<0.05, ** P<0.01, *** P<0.001). (b) Immunoblot analysis of the lines shown in (a) probed with α-*At*GRP7 and α-*At*GRP8 antipeptide antibodies. Amidoblack staining of the membrane (AB) served as loading control. The uncropped blot is shown in Suppl. Fig. S1.

Together, these data suggest that exchange of R141 to either alanine or lysine is not critical for the floral promotive effect of *At*GRP7 or the negative impact on *At*GRP8.

In parallel, we complemented the *grp7-1* mutant with constructs expressing the *At*GRP7-GFP fusion protein under control of the endogenous promoter and 3’ untranslated region as well as constructs expressing *At*GRP7-GFP with the R141K or R141A mutation. The *grp7-1* mutant complemented with wt *At*GRP7-GFP flowered with significantly fewer leaves than the *grp7-1* mutant. Three independent lines expressing *At*GRP7-GFP R141A and three independent lines expressing *At*GRP7-GFP R141K flowered with a similar leaf number as the line complemented with wt *At*GRP7-GFP, again indicating that R141 is not critical for the floral promotive effect (Fig. 2a). Protein levels for *At*GRP7-GFP R141A were higher than for *At*GRP7-GFP R141K, but only line 38 showed *At*GRP8 levels like wt whereas in lines 48 and 50 *At*GRP8 was higher than wt. The two *At*GRP7-GFP R141K lines 4 and 50 with higher *At*GRP7 levels had *At*GRP8 levels like wt, while the line 23 with the lowest *At*GRP7-GFP level showed a strong signal for *At*GRP8 (Fig. 2b). This was also the line with the highest leaf number at bolting, again indicating a certain dose-dependency of the *At*GRP7-GFP protein abundance and flowering phenotype. All in all, *At*GRP7-GFP complemented the *grp7-1* mutant similar to the wt genomic fragment, without further impact of the R141 mutational status.

**Figure 2.**
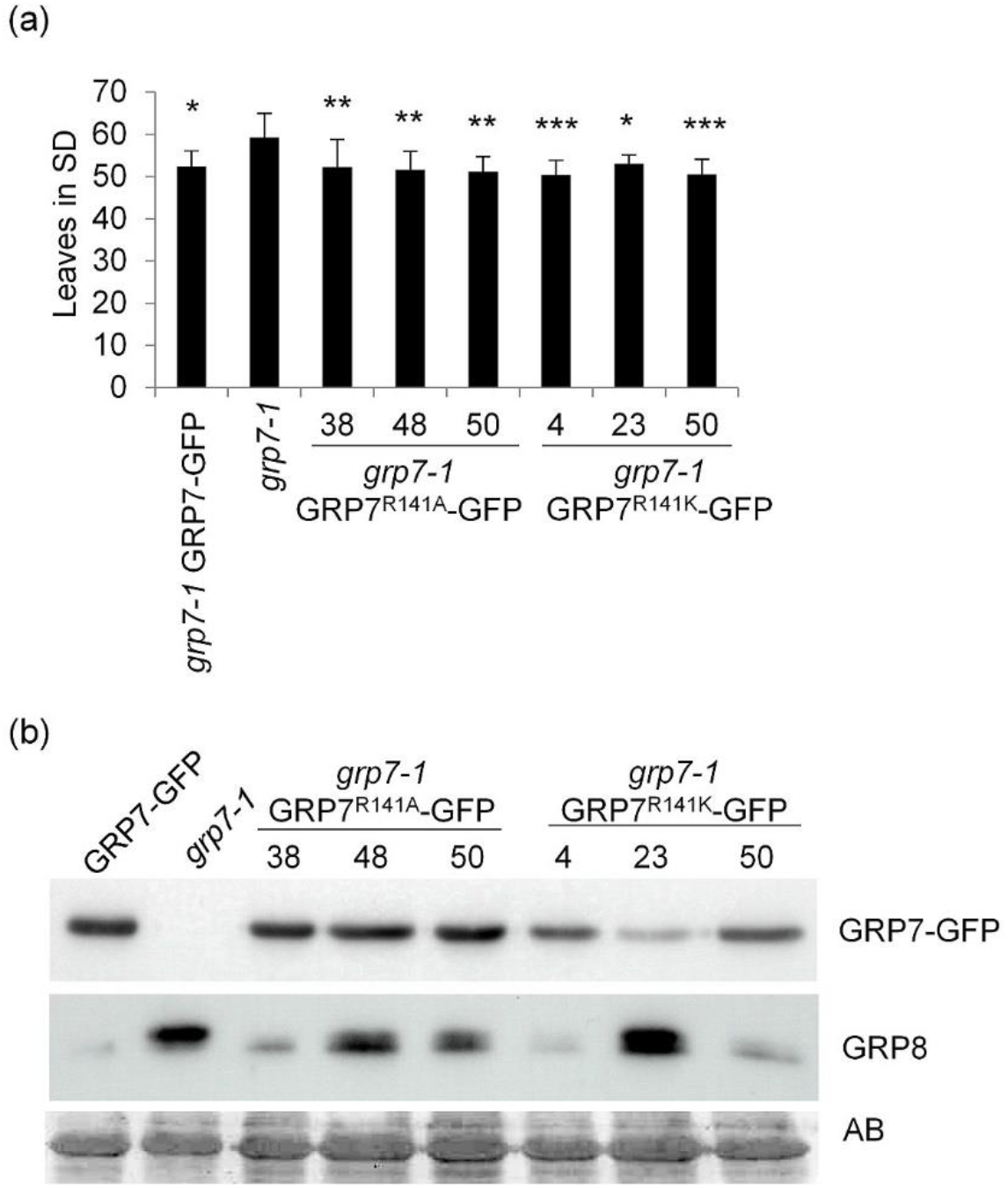
Flowering time of *grp7-1* lines complemented with *At*GRP7:: *At*GRP7-GFP and R141 variants. (a) The *grp7*-1mutant and the *grp7-1* line complemented with *At*GRP7:: *At*GRP7-GFP, *At*GRP7:: *At*GRP7 R141A-GFP, and *At*GRP7:: *At*GRP7 R14KA -GFP were grown in SDs (n = 15-20). The number of rosette leaves are shown as mean ± SD. ANOVA followed by a Dunnett’s test was performed to determine statistical significance (* P<0.05, ** P<0.01, *** P<0.001). (b) Immunoblot analysis of the lines shown in (a) probed with α-GFP antibody and α-GRP8 antipeptide antibody. Amidoblack (AB) staining served as loading control. The uncropped blot is shown in Suppl. Fig. S2.

### 2.2 Overexpression of *At*GRP7 leads to dose-dependent early flowering independent of R141 mutations

Constitutive overexpression of *At*GRP7 causes plants to flower earlier than wt plants (24,25). We introduced the R414K and R141A mutations into the *AtGRP7* cDNA driven by the CaMV promoter. In addition, we generated an R141F mutation to mimic constitutive R141 methylation.

Two independent lines overexpressing wt *At*GRP7 flowered with less leaves than Col-0 and had *At*GRP8 levels below the detection limit of the antibody, due to negative regulation of *AtGRP8* by *At*GRP7 (26). Seven lines overexpressing *At*GRP7 R141A coming from four independent transformation events flowered with less leaves than Col-0 and not significantly different from plants overexpressing the authentic *At*GRP7 protein. The *At*GRP7 R141A level was variable in these lines. While lines 1.1, 1.3, 7, 15.2 and 18 had high levels of *At*GRP7 and barely detectable levels of *At*GRP8, lines 1.2 and 15.1 expressed *At*GRP7 R141A to a similar extent as wt plants. Intriguingly, line 1.2 had no detectable *At*GRP8, while in line 15.1 the level was comparable to Col-0 (Fig. 3a, b).

**Figure 3.**
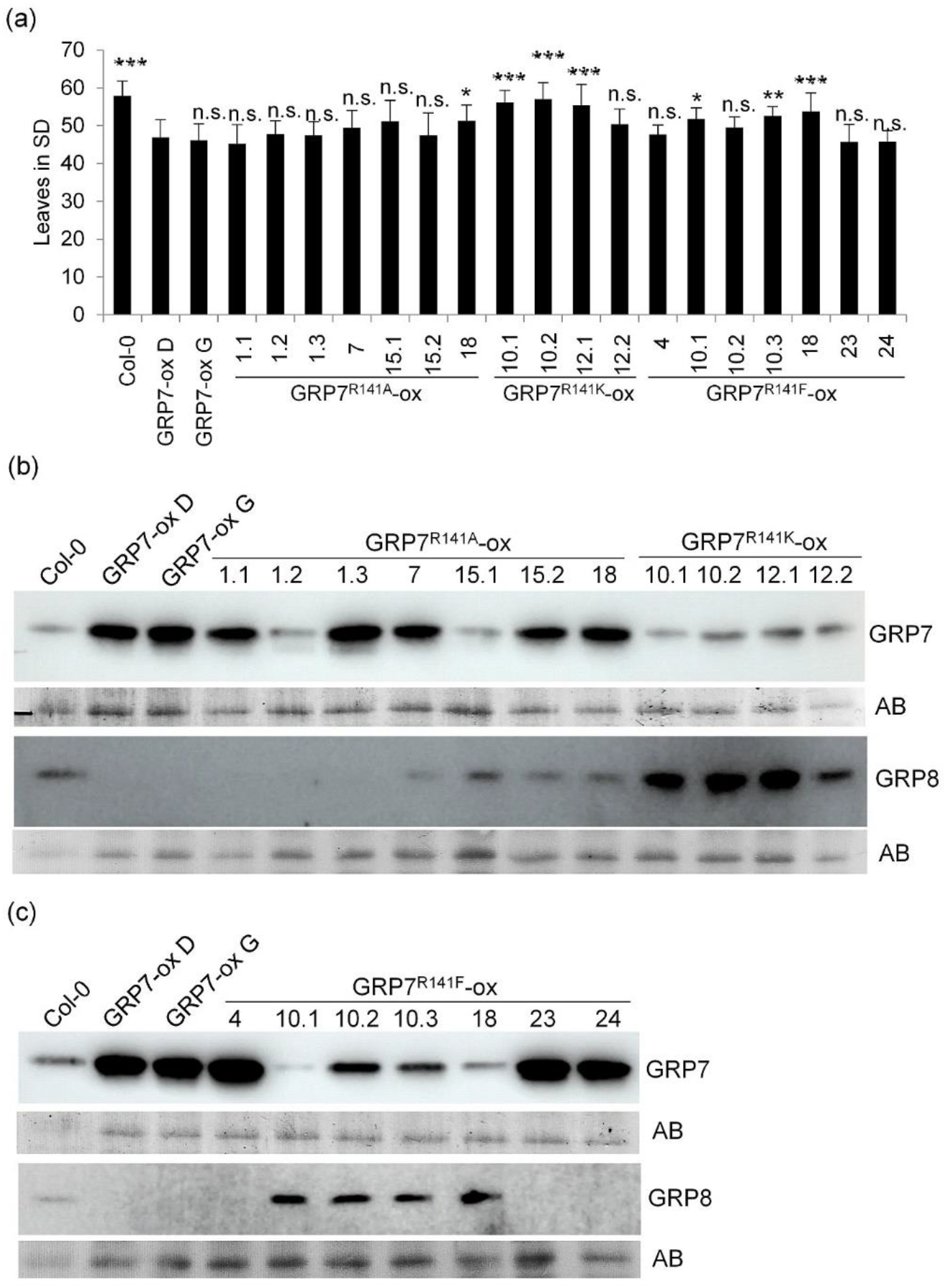
Flowering time of plants overexpressing *At*GRP7, *At*GRP7 R141A, *At*GRP7 R141K and *At*GRP7 R141F. (a) *At*GRP7-ox D and plant lines overexpressing *At*GRP7 R141A, *At*GRP7 R141K and *At*GRP7 R141F were grown in SDs (n = 15-20). The number of rosette leaves are shown as mean ± SD. ANOVA followed by a Dunnett’s test was performed to determine statistical significance (* P<0.05, ** P<0.01, *** P<0.001, n.s., not significant). (b) Immunoblot analysis of *At*GRP7 R141A-ox, *At*GRP7 R141K-ox and (c) *At*GRP7 R141F-ox with α-GRP7 and α-GRP8 antipeptide antibodies. Amidoblack (AB) staining served as loading control. The uncropped blot is shown in Suppl. Fig. S3.

From four lines with the *At*GRP7 R141K overexpression construct only line 12.2 flowered like plants overexpressing wild type *At*GRP7 and three lines flowered as Col-0. All lines did not overexpress *At*GRP7 R141K, but had protein levels comparable to Col-0 instead. *At*GRP8 levels were higher than wt in all lines, indicating that the R141K exchange might attenuate the downregulating effect from *At*GRP7-ox on *At*GRP8. (Fig. 3 a, b).

Four independent lines overexpressing *At*GRP7 R141F (4, 10.2, 23, 24) also flowered with a similar leaf number as plants overexpressing the authentic *At*GRP7 protein. These lines also had the highest *At*GRP7 protein levels. Three lines with lower *At*GRP7 levels flowered with leaf numbers intermediate between Col-0 and *At*GRP7-ox D. Again, *At*GRP8 levels were reduced below the detection limit in lines with high *At*GRP7 R141F expression, and somewhat higher than Col-0 in lines with Col-0-like levels of *At*GRP7 R141F (Fig. 3 c). Taken together, our data suggest that *At*GRP7 reduced the number of leaves at bolting irrespective of R141 mutations. Instead, flowering behaviour of the different lines was more dependent on the overall expression level of the different *At*GRP7 R141 variants.

### 2.3 *At*GRP7 R141A attenuates the response to abscisic acid

As we did not observe strong effects of *At*GRP7 R141 mutations on flowering time, we set out to find a different readout for a phenotypic impact of R141 mutations. In *prmt5-5* plants with impaired methylation, alternative splicing events in genes associated to abiotic stress are overrepresented indicating a strong effect of arginine methylation on stress response (27). One of the earliest reactions to abiotic stress is the accumulation of abscisic acid (ABA) that inhibits germination and other developmental processes to help the plant cope with unfavorable conditions. We therefore monitored germination under ABA treatment. Wt and *grp7-1* similarly repressed germination after 2 days of 1 μM ABA treatment so that only 30%-40% of seeds germinated. *At*GRP7-ox D was almost insensitive to ABA and germination was around 90%. The *At*GRP7 R141A overexpressing line 1.1 showed significantly less inhibition of germination than wt, but did not reach the level of *At*GRP7-ox D. In line 15.2, inhibition was even weaker so that it was not statistically different from wt any more (Fig. 4a). Immediately after germination, we also monitored the fully open cotyledon stage (“greening”). Again, no significant difference was visible between wt and *grp7-1*, with ∼30% of the plants reaching this stage. Still, a higher percentage of seedlings of *At*GRP7ox D reached the fully open stage compared to wt, while greening was inhibited more strongly in *At*GRP7 R141A ox 1.1 and 15.2 compared to the plants that express the wild type version (Fig. 4 b). This indicates that overexpression of *At*GRP7 leads to reduced ABA sensitivity and that the R141A mutation impacts the ABA related phenotypes germination and greening, pointing towards a dedicated role for R141 and, by extension, its methylation under stress. This is in concordance with an enhanced sensitivity to ABA observed in *prmt5-5* (7,15).

**Figure 4.**
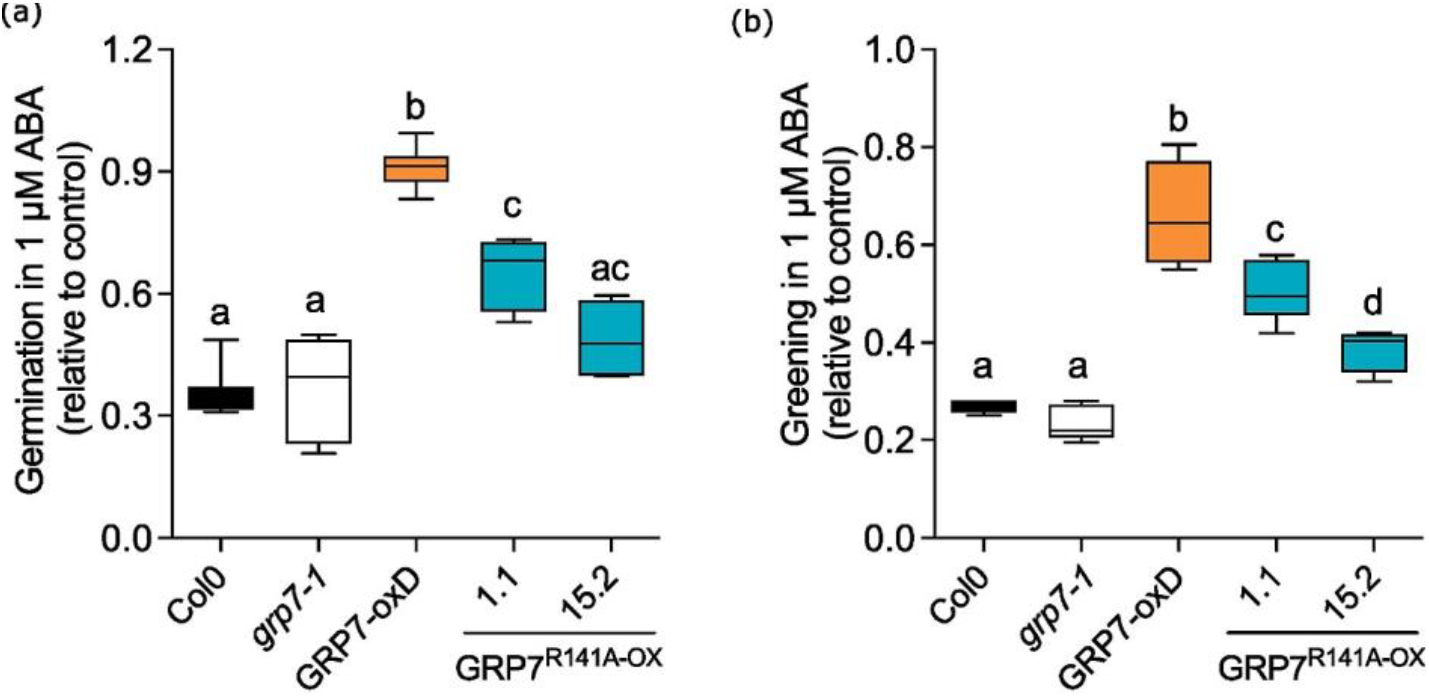
GRP7 R141 is required for the response to ABA stress. Col-0, *grp7-1, At*GRP7-ox D and *At*GRP7 R141A ox 1.1 were grown in 16h light/8h dark on ½ MS plates containing 1 μM ABA. Control plants were grown without ABA. (a) Germination was scored two days after start of the experiment. (b) Fully expanded cotyledons were scored after 7 days. Data are mean ± SD of six independent biological replicates (100 seeds per experiment). One way ANOVA was performed to assess statistical significance. Different letters indicate a significant difference at p≤0.05 (Tukey test).

### 2.4 *In vivo* dimethylation status of *At*GRP7

Next, we addressed the *in vivo* methylation status of *At*GRP7 under 12h light/12h dark conditions with the SYM10 antibody specifically detecting symmetrically dimethylated arginine. To distinguish between *At*GRP7 and other dimethylated proteins, we grew plants expressing *At*GRP7-GFP in the *grp7-1* background and plants expressing *At*GRP7-GFP in *prmt5-1* and *prmt5-5* backgrounds and pulled down *At*GRP7-GFP from total protein extracts with GFP-Trap beads. Detection with an antibody against GFP detected the fusion protein in the input and proved successful pulldown of the fusion protein in the immunoprecipitation (IP) in all three backgrounds, but indicated that the expression was lower in the *prmt5-1* and *prmt5-5* backgrounds (Fig. 5a). As expected, no *At*GRP7-GFP was pulled down from *prmt5-1* and *prmt5-5* alone. Detection with the SYM10 antibody showed multiple bands of potentially dimethylated proteins in the *At*GRP7-GFP background. A similar band pattern was also observed in *prmt5* mutant backgrounds with one protein band at around 55 kDa missing and another one around 15 kDa strongly reduced (Fig. 5a). The IP proved that *At*GRP7-GFP was dimethylated in the *grp7-1* background where PRMT5 is active, but a weak signal was also detected in *prmt5-1* GRP7-GFP and *prmt5-5* GRP7-GFP (Fig. 5a). Amidoblack staining of the membrane proved that more protein was pulled down from *At*GRP7-GFP in *grp7-1* than from the *prmt5-1* and *prmt5-5* backgrounds, reflecting the difference in signal strength from the SYM10 antibody, as previously observed (7) (Fig. 5a). To independently verify this, we independently repeated the IP experiment, this time including wt plants. Detection with GFP again proved successful IP of *At*GRP7-GFP (Fig. 5b). To detect the dimethylation status, we used the αsdmR antibody (Cell Signaling Technologies). The input showed a band larger than 55 kDa in Col-0 and plants expressing *At*GRP7-GFP. Interestingly, in *At*GRP7-GFP and *At*GRP7 GFP in *prmt5-1* and *prmt5-5* a signal around 40 kDa was also present, likely belonging to *At*GRP7-GFP (Fig. 5b). The IP with the αsdmR antibody confirmed *At*GRP7-GFP as dimethylated in *At*GRP7-GFP, *prmt5-1 At*GRP7-GFP and *prmt5-5 At*GRP7-GFP (Fig.5b). Again, the signal was weaker in the *prmt5* mutant backgrounds, but amidoblack staining indicated that also less protein was pulled down from these lines (Fig. 5b). Taken together, our results indicate that *At*GRP7-GFP is still methylated in *prmt5-1* and *prmt5-5*, but possibly at a somewhat reduced level. This would hint that PRMT5 might not be the only methyltransferase responsible for symmetric dimethylation of *At*GRP7.

**Figure 5.**
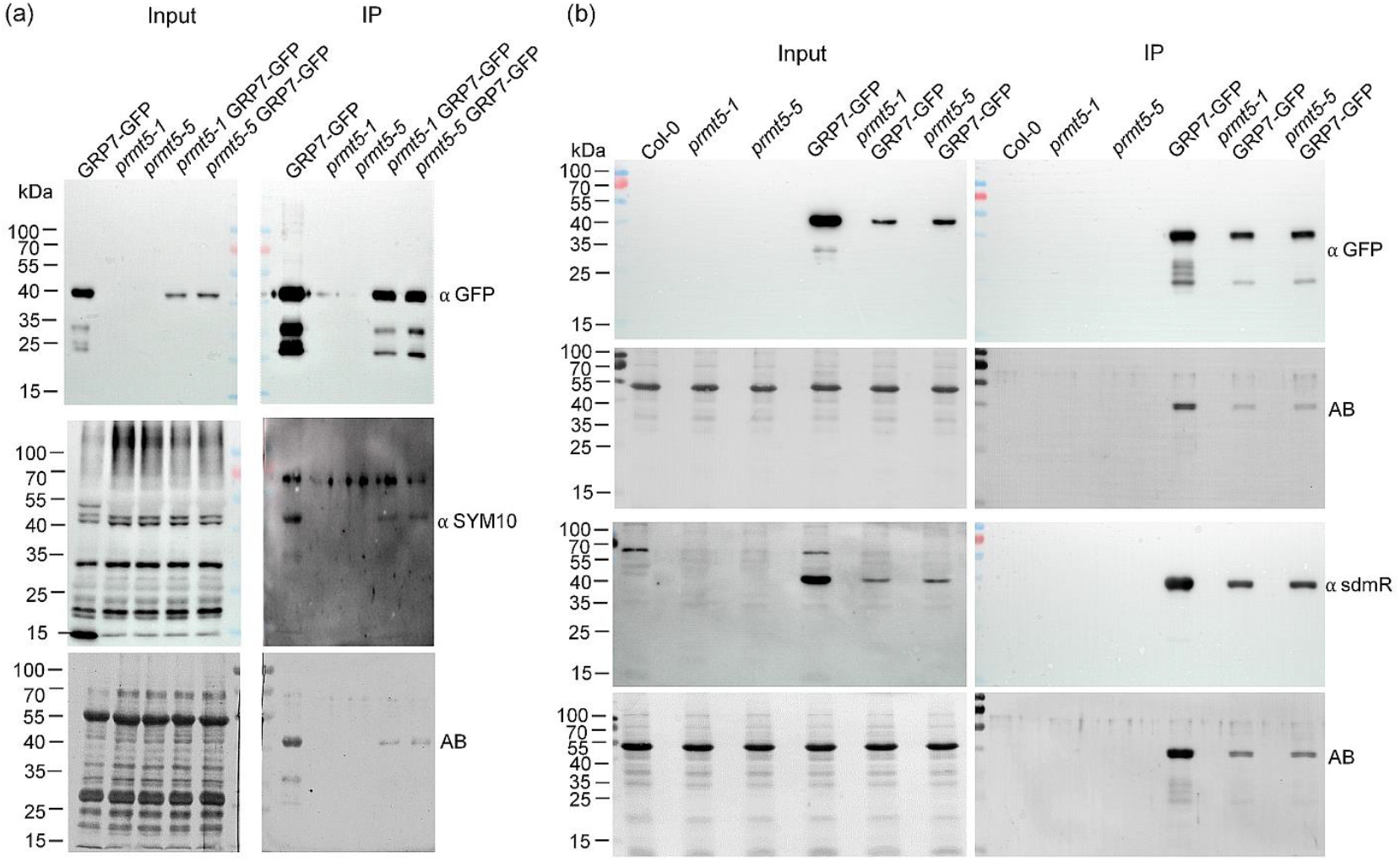
*In vivo* arginine dimethylation status of *At*GRP7-GFP (a) *At*GRP7-GFP, *prmt5-1, prmt5-5* and *At*GRP7-GFP in *prmt5-1* and *prmt5-5* were grown in 12h light/12h dark on ½ MS plates. Native protein extracts were subjected to immunoprecipitation with GFP-Trap beads to enrich *At*GRP7-GFP and membranes were probed with antibodies against GFP and the SYM10 antibody. Amidoblack (AB) staining served as loading control. (b) Col-0, *prmt5-1, prmt5-5, At*GRP7-GFP and *At*GRP7-GFP in *prmt5-1* and *prmt5-5* were grown in 12h light/12h dark on ½ MS plates. Native protein extracts were subjected to immunoprecipitation with GFP-Trap beads to enrich *At*GRP7-GFP and membranes were probed with antibodies against GFP and symmetrical dimethylated arginine residues (sdmR). Amidoblack (AB) staining served as loading control. The uncropped blots are shown in Suppl. Fig. S4.

## 3. Discussion

The role of the arginine methyltransferase PRMT5 in regulating various biological processes, including flowering time in Arabidopsis, has become increasingly evident through recent studies. PRMT5 catalyzes symmetric dimethylation of arginine residues, a modification crucial for controlling gene expression, splicing, and protein-protein interactions (1,12,13). Use of antibodies against symmetric dimethylarginine has unraveled a defined subset of proteins modified by PRMT5, among them core spliceosomal components and other RNA-binding proteins (10). In the context of flowering, PRMT5 appears to regulate this process at least partly through its effect on AS of *FLK*, ultimately leading to altered levels of the key floral repressor FLC. The glycine-rich RNA-binding protein *At*GRP7, which is involved in circadian rhythm and RNA processing, was shown to be modified by PRTM5 on R141 located in the glycine-rich stretch C-terminal stretch. Here, we addressed the question of whether *At*GRP7 arginine dimethylation on R141 is essential for its role in flowering time regulation. Our results demonstrate that substituting R141 with alanine (R141A) or lysine (R141K) did not critically impair the floral promoting effect of *At*GRP7. Transgenic lines expressing either mutated version of *At*GRP7 (R141A or R141K) flowered similarly to wt plants when reintroduced into the *grp7-1* mutant background. This suggests that R141 methylation in the glycine-rich C-terminus is not essential for flowering promotion.

Further supporting this, overexpression studies showed that plants overexpressing *At*GRP7 with these mutations still flowered earlier than wild-type plants, similar to those overexpressing the unmodified *At*GRP7. This indicates that the overall expression level of *At*GRP7, rather than the methylation status of R141, is the primary determinant of its role in promoting flowering.

However, the repressive effect of *At*GRP7 on *At*GRP8 expression appeared to be somewhat dependent on the mutation status of R141. Lines expressing the R141K mutation displayed higher *At*GRP8 levels compared to wt, indicating a potential reduction in the repressive capacity of *At*GRP7 when this residue is altered. This points to a nuanced role for R141 methylation, possibly influencing specific protein interactions rather than the broad regulatory functions of *At*GRP7.

Moreover, investigation of the *in vivo* methylation status of *At*GRP7 in *prmt5* mutant backgrounds provided intriguing insights. Although PRMT5 was shown to be responsible for *At*GRP7 and *At*GRP8 methylation, both were not detected as dimethylated any more in the *prmt5* mutant and R141 was identified as the residue modified (10), here dimethylation of *At*GRP7 was still observed, albeit at reduced levels, in *prmt5* mutants. This implies a potential redundancy in the methylation machinery within Arabidopsis where other methyltransferases compensate for the loss of PRMT5.

A recent study aiming to uncover the Arabidopsis methylome identified 236 arginine methylation sites on 149 non-histone proteins. Of those, only 22 proteins contained 29 different symmetric dimethylation sites (28). While *At*GRP8 R141 was identified as symmetrical dimethylated, *At*GRP7 was not identified in this study. Instead, *At*GRP7 R141 appeared as monomethylated. These contradictory findings may hint that arginine methylation is a highly dynamic process, largely influenced by environmental conditions or stress. Additionally, other R residues could be methylated as well, although R47 and R96 were also only detected as monomethylated (29). Interestingly, it was reported recently that the commonly used tryptic digest of proteins for mass spectrometry leads short and highly polar peptides that are difficult to separate and sequence, leading to insufficient coverage of peptides in arginine rich regions (30). This may explain some of the conflicting data in the literature on the methylation status of the peptides derived from the intrinsically disordered C-terminus of *At*GRP7 that is enriched with interspersed arginine residues including R141 (23,31,32).

In summary, while mutations of *At*GRP7 R141 to alanine, lysine or phenylalanine are not critical for its floral promotive function, they may influence specific interactions that affect other regulatory pathways, such as the repression of *AtGRP8*, and sensitivity to ABA. These findings highlight the complexity of post-translational modifications in plant development and suggest that the role of PRMT5 in flowering is multifaceted, possibly involving additional targets and compensatory mechanisms.

## 4. Materials and Methods

### 4.1. Constructs and transgenic plants

The *At*GRP7::*At*GRP7-GFP line expressing *At*GRP7::GFP under control of the *At*GRP7 promoter has been described (33,34). The *grp7-1* 8i line has an RNAi construct against *At*GRP8 to counteract elevated *At*GRP8 level due to relief of repression by *At*GRP7 in *grp7-1* (24). *At*GRP7-ox plants express the *At*GRP7 coding sequence under control of the Cauliflower Mosaic Virus (CaMV) 35S promoter (35).

Gene fragment encoding the glycine-rich C-terminal part of *At*GRP7 was synthesized with arginine 141 mutated to alanine (R141A) or to lysine (R141K) (Eurofins).

To overexpress the mutant proteins under control of the double CaMV promoter, the corresponding fragments in pRT103-*At*GRP7 were replaced by the R141A or R141K mutant fragments via XcmI-BamHI digest. To overexpress the *At*GRP7 variant where R141 was mutated to phenylalanine (R141F) site-directed mutagenesis was performed on pRT-*At*GRP7 R141A with primers indicated in Suppl. Table S1.

To generate mutant *At*GRP7-GFP fusion proteins, a 170 nucleotide XcmI-BbsI fragment of *At*GRP7::*At*GRP7-GFP was replaced by the R141A or R141K mutant fragments via XcmI-BbsI digest to yield pRT-GRP7::GRP7-R141A-GFP and pRT-pGRP7::GRP7-R141K-GFP constructs.

To obtain genomic clones with the R141 mutations, the BbsI-XcmI fragments were cloned into a 3 kb genomic *At*GRP7 fragment including 1.4 kb of the *At*GRP7 promoter (33)

All constructs were verified by sequencing and the expression cassettes were mobilized to the binary vector HPT1 and introduced into *Arabidopsis thaliana* via Agrobacterium-mediated floral dip.

*At*GRP7-GFP plants were crossed to *prmt5-1* and *prmt5-5* to obtain *prmt5-1 At*GRP7-GFP and *prmt5-5 At*GRP7-GFP. Homozygous plants were identified in the F2 generation based on GFP fluorescence, immunoblot against *At*GRP7-GFP and PCR-genotyping of the respective *prmt5* alleles with primers indicated in Suppl. Table S1.

### 4.2. Determination of flowering time

Seeds were sown on soil, stratified at 4 °C for two days, and germinated and grown in SDs (8-hr light/16-hr dark cycles) or LDs (16-hr light/8-hr dark cycles). Plants were grown in a randomized fashion at 20 °C in Percival incubators AR66-L3 (CLF Laboratories). Flowering time was determined by counting the rosette leaves once the bolt was 0.5 cm tall. Mean values ± SD were calculated (25). For statistical analysis, ANOVA followed by a Dunnett test were performed in case of a normal distribution of the data. Otherwise, a Kruskal-Wallis test was performed.

### 4.3. Physiological response to ABA

Seeds were sown on MS medium supplemented with 1 μM ABA. The proportion of germinated seeds was scored after 48 h while greening was scored after 7 days. Both parameters correspond to radicle emergence and fully opened cotyledons, 0.5 and 1 stages, respectively, according to Boyes et al. (36). Approximately, 100 seeds were processed per line in each experiment. The data were subjected to way ANOVA and post hoc comparisons were performed with Tukey’s multiple range test.

### 4.4. Immunoblot analysis

Protein extracts were prepared as previously described (37). Western blot analysis with anti-peptide antibodies against *At*GRP7 and *At*GRP8 was done as described (38). Amidoblack staining of the membrane served as loading control.

For the Immunoprecipitation to detect symmetrical dimethylated arginine, protein extracts were prepared in native buffer (50 mM Tris-HCl, ph 7.5, 100 mM NaCl, 10% (v/v) glycerol, Complete™ Protease inhibitor cocktail tablet EDTAfree (Roche), 2mM PMSF). GFP-tagged proteins were pulled down using GFP Trap beads (ChromoTek & Proteintech Germany). After four washing steps with IP wash-buffer (50 mM Tris-HCl pH 7.5, 100 mM NaCl, 10% (v/v) glycerol, 0.05% (v/v) Triton X-100), the beads were boiled in Laemmli buffer and directly loaded onto SDS-PAGE gels. Symmetrical dimethylated *At*GRP7-GFP was detected with SYM10 antibody (Sigma-Aldrich, No. 07-412) or αsdmR antibody (Cell Signaling Technologies, No. 13222S). *At*GRP7-GFP was detected with αGFP coupled to horseradish peroxidase (Miltenyi Biotec, No. 130-091-833).

## Supplementary Materials

The following supporting information can be downloaded at: www.mdpi.com/xxx/s1, Figure S1: Uncropped blots corresponding to Fig.1; Figure S2: Uncropped blots corresponding to Fig.2; Figure S3: Uncropped blots corresponding to Fig.3; Figure S4: Uncropped blots corresponding to Fig.5; Table S1: Primers used in this study

## Author Contributions

Conceptualization, J.L.M. and D.S.; methodology, A.S., J.L.M. and D.S.; investigation, A.S, K.D., C.N., M.J.I., M.J.L.; writing—original draft preparation, A.S. and D.S.; writing—review and editing, A.S., J.L.M. and D.S..; supervision, D.S.; project administration, J.L.M. and D.S.; funding acquisition, D.S., J.L.M. All authors have read and agreed to the published version of the manuscript.

## Data Availability Statement

The original contributions presented in the study are included in the article/supplementary material, further inquiries can be directed to the corresponding author.

## Acknowledgments

We thank Elisabeth Klemme, Kristina Neudorf, and Frederik Dombert for expert technical assistance.

